# 3D-Feedy: An open and portable membrane feeder for mosquito research

**DOI:** 10.1101/2025.10.28.685147

**Authors:** Gleb Ens, Ariel Roselin Lewin, Doron SY Zaada, Josef Doron, Ziv Gilan, Ariel Livne, Elly Ordan, Philippos Aris Papathanos

## Abstract

Female mosquitoes must feed on blood to produce eggs, a natural behavior that also drives disease transmission. Vector biology research and the development of novel control tools depend on reliable methods for blood feeding of laboratory-reared mosquitoes. Artificial membrane feeders have provided a practical alternative to live animals for decades. However, commercial systems are typically expensive, difficult to repair and are tethered to external water baths or bench-top power supplies. Their limited capacity, usually only a few cages at once, further restricts their use, making them impractical in field settings or large insectaries where dozens of experimental crosses must be fed daily.

Here we describe 3D-Feedy, an open-hardware, inexpensive membrane feeder that delivers electrically heated membrane feeding using simple, readily available parts. Powered via USB-C, including smartphone powerbanks, it is portable and straightforward to assemble with only basic tools and skills. In direct comparison with the widely used Hemotek feeder in *Anopheles gambiae* and *Aedes albopictus*, the 3D-Feedy achieved equal or higher feeding rates and significantly greater egg production while maintaining comparable larval hatching rates. These results validate 3D-Feedy as a practical and accessible tool for mosquito rearing and experimental studies. All design files and instructions are provided as open hardware resources.

## Introduction

Blood feeding is essential for maintaining mosquito colonies and for experimental research in vector biology. Female mosquitoes require a blood meal to initiate egg production, and although live animals have historically been used for this purpose, such methods present regulatory, ethical, and logistical challenges. Artificial membrane feeders address these limitations by delivering heated blood meals in a controlled and reproducible manner. In some applications, such as feeding with infected blood, they are indispensable (Dong *et al*., 2011; Pascini *et al*., 2022; Graumans *et al*., 2023; Li *et al*., 2025).

Two main designs dominate commercially available systems: water-jacketed glass feeders, first introduced by Rutledge et al. in the 1960s (Rutledge, Ward and Gould, 1964) and electrically heated systems such as Hemotek (Hemotek Ltd., Blackburn, UK). Both are reliable, but they are expensive, dependent on specialized suppliers, and are difficult to repair. They also lack portability, requiring cages to be brought to feeding stations near water baths or bench-top power supplies, which limits use in insectaries and field settings. Finally, both designs typically accommodate only four to six cages at once; in large insectaries where dozens of cages can be fed daily, this constraint translates into substantial time and labor cycling cages through a small number of stations.

Several groups have introduced designs aimed at lowering costs and broadening accessibility. Many rely on passive heating, where the membrane is warmed by contact with a pre-heated thermal reservoir, e.g. an insulated cup, water-filled vessel, metal block (Tseng, 2003; Costa-da-Silva et al., 2013; Finlayson, Saingamsook and Somboon, 2015; Siria et al., 2018; Sri-in et al., 2020). However, without active heating, the reservoir cools quickly once removed from the heat source, narrowing the feeding window and forcing repeated reheating, an issue that becomes acute when feeding multiple cages.

More recent approaches have incorporated 3D-printed components to replace glass feeders and simplify assembly (Witmer *et al*., 2018; Graumans *et al*., 2020). These reduce dependence on specialized fabrication but most still rely on external circulating water baths for heating, limiting portability and constraining their use outside a typical laboratory setting. Dewsnup et al. recently described the SLAM feeder, an open-hardware electrically heated alternative that demonstrated how membrane feeders can be made accessible through do-it-yourself (DIY) construction (Dewsnup *et al*., 2023).

Here we introduce 3D-Feedy, an open-hardware device assembled from widely available electronic components and machined or 3D-printed parts. It offers the reliability of electrically heated systems while remaining inexpensive, portable, and straightforward to assemble and repair. Powered via standard USB-C, including from common powerbanks, it can be used in insectaries, within incubators, and at field sites. Assembly requires only basic tools and inexpensive materials, can be completed in a few hours for approximately 30 USD, and is simple enough to serve as a first electronics project. We benchmark the performance of our 3D-Feedy against the gold-standard Hemotek system in *An. gambiae* and *Ae. albopictus*, comparing feeding success, fecundity, and egg viability. Our results validate the 3D-Feedy as a practical and accessible alternative for mosquito research.

## Results

### Design and operation of the 3D-Feedy membrane feeder

The 3D-Feedy is constructed from a small number of inexpensive, readily-available components (**Figure 1A–D**). At its core is a compact W1209 digital temperature controller that regulates a resistive film heater bonded to the back of a machined aluminum block (44 × 55 × 5 mm), referred to as the heated block. A cavity within the heated block houses the digital temperature probe, which provides continuous feedback to the thermostat. Power is supplied via a USB-C breakout board, enabling operation from standard 5 V powerbanks or wall chargers. All electronic components are enclosed within a custom 3D-printed housing designed to secure the heated block, protect circuitry, and allow simple disassembly for maintenance or replacement (**Figure 1A**). The assembled device is lightweight and compact, designed to rest directly on top of mosquito cages during feeding (**Figure 1B**). Blood is presented to mosquitoes on a second, larger (100 × 100 × 5 mm, rounded corners) aluminum block termed the feeding plate. The rear of the plate is machined to create three recesses: a central pocket that exactly accommodates the 44 x 55 mm heated block and two parallel grooves that align and lock the feeding plate into position using 3D-printed alignment tabs. During feeding, the feeding plate is wrapped on its front side with a Parafilm membrane (8 x 10 cm) forming a pocket on its surface. 2 ml of thawed blood are pipetted into the Parafilm membrane of the feeding plate, which is then placed beneath the feeder (**Figure 1C**).

**Figure 1.**
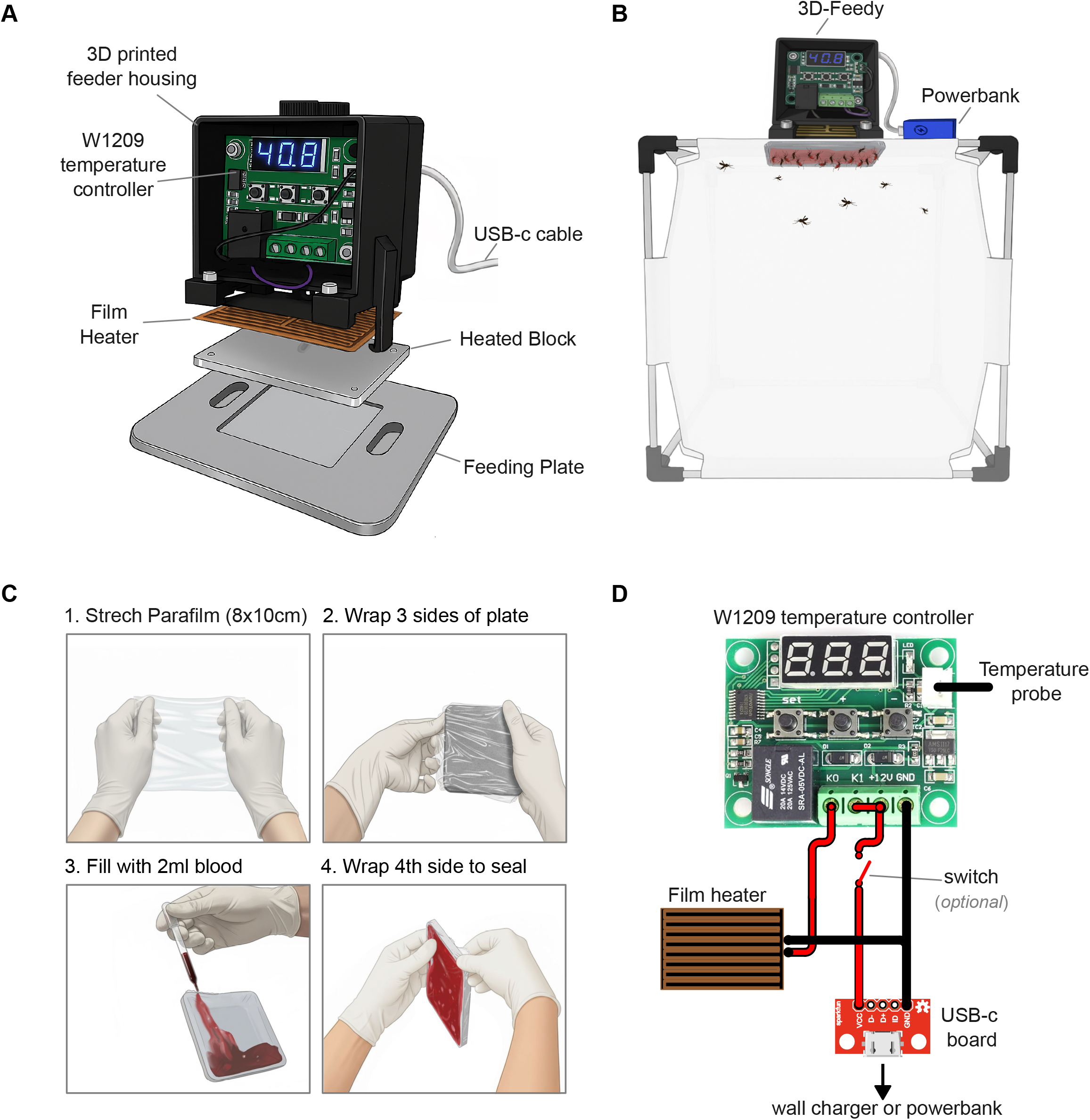
Design, use, and wiring of the 3D-Feedy membrane feeder. **(A)** Exploded view showing the components of the 3D-Feedy. **(B)** Drawing of an assembled feeder in use on top of a mosquito cage, powered by a USB-C powerbank (in blue). **(C)** Step-by-step preparation of the feeding plate. **(D)** Wiring schematic of the 3D-Feedy system showing connections between the USB-C power input, W1209 temperature controller, resistive film heater, and temperature probe.

Calibration of the thermostat setpoint is required once during initial assembly to ensure accurate feeding temperature. This is done by measuring the surface temperature of the feeding plate with an independent thermometer, and then adjusting the thermostat until the mosquito-facing surface reaches 37□°C. In practice, setting the thermostat to 40□°C yields this plate temperature. Calibration needs to be performed only once, as the setting is stored in memory.

The wiring layout is simple **(Figure 1D)** with the USB-C power input connected to the W1209 temperature controller, which in turn regulates the resistive film heater and receives feedback from the temperature probe. This minimal wiring design allows rapid assembly and easy replacement of individual components. All design files are provided as STL and STEP models for 3D-printing and machining (Supplementary Files 1–4). In operation, the feeder connects directly to any USB-C power source. A 5000 mAh powerbank can power the feeder continuously for around 5-6 hours, allowing multiple cages to be fed sequentially.

### Benchmarking mosquito blood feeding

We benchmarked the 3D-Feedy membrane feeder against the commercial Hemotek system (Discovery Workshops, Accrington, UK) system by assessing female feeding rates, egg production, and hatching success in *An. gambiae* and *Ae. albopictus* (Figure 2). For each feeder type and species, three replicate cages of 50 mated females were given access to a blood feeder for 30 minutes, after which the proportion of females that successfully engorged was recorded. From each cage, three independent groups of five fed females were isolated for downstream reproductive assays, yielding nine subgroups per treatment. Egg production was quantified per female, and hatching success was calculated as the percentage of larvae emerging from the total number of eggs laid (**Figure 2**).

**Figure 2.**
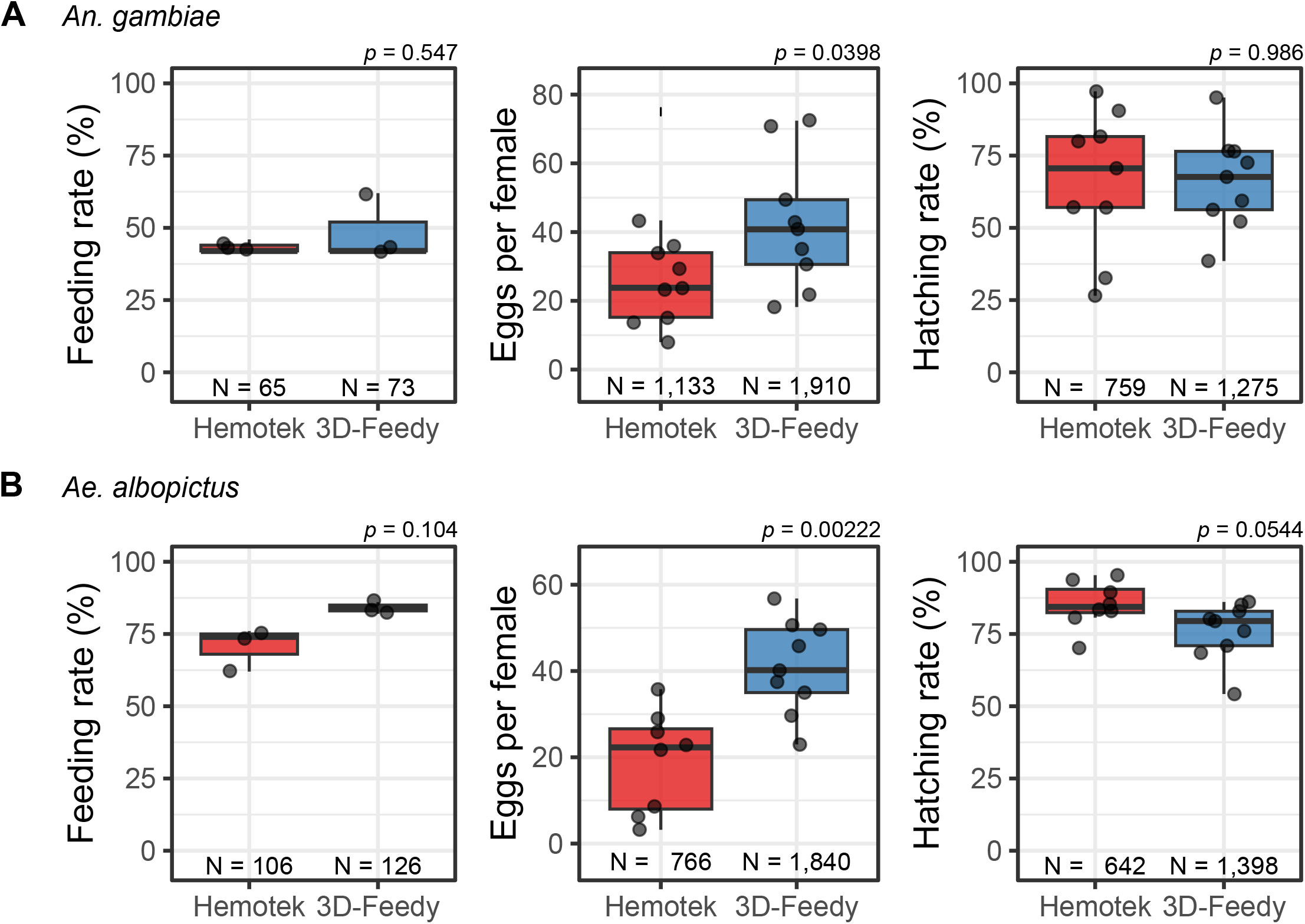
Comparison of mosquito feeding performance between the 3D-Feedy and Hemotek systems. Percentage of fed females (feeding rate), eggs per female, and hatching rate of *Anopheles gambiae* **(A)** and *Aedes albopictus* **(B)** females fed with the Hemotek or 3D-Feedy systems. Bars show means ± SD, with individual data points representing biological replicates. Statistical significance was determined using two-sample t-tests or Wilcoxon rank-sum tests as appropriate. Numbers below each plot indicate the total number of individuals scored in each experiment across all replicates.

For *An. gambiae*, feeding rates were similar between feeders with 48.7 ± 6.7% of females feeding from the 3D-Feedy compared to 43.3 ± 1.3% from Hemotek (Wilcoxon rank-sum test, *W* = 5, *p* = 1.00). However, 3D-Feedy-fed females laid significantly more eggs per female than those fed with Hemotek (42.44 ± 25.18 vs 25.18 ± 13.13; *t* = 2.31, *df* = 16, *p* = 0.035). Egg hatching rates were nearly identical between the two feeders (0.66 ± 0.07 vs 0.66 ± 0.08; *t* = 0.017, *df* = 16, *p* = 0.99; **Figure 2A**). For *Ae. albopictus*, females fed with the 3D-Feedy system achieved a higher feeding rate than those fed with Hemotek (0.84 ± 0.05 vs 0.71 ± 0.06; *t* = 2.95, *df* = 4, *p* = 0.042). Egg production was significantly higher for females fed with the 3D-Feedy system, (40.89 ± 19.15 vs 19.15 ± 14.38; *t* = 3.96, *df* = 15, *p* = 0.001), corresponding to more than a twofold increase. Egg hatching rates were modestly lower for 3D-Feedy-fed females (0.76 ± 0.08) than in Hemotek (0.85 ± 0.06), but the difference was not statistically significant (*t* = −2.06, *df* = 15, *p* = 0.057; **Figure 2B**).

A two-way ANOVA combining both species revealed a strong species effect on feeding rate (*F*□,□ = 58.91, *p* < 0.001) and a near-significant effect of feeding system (*F*□,□ = 5.23, *p* = 0.052). For egg production, the interaction between species and feeding system was significant (*F*□,□ = 6.95, *p* = 0.030), indicating that the benefit of the 3D-Feedy system differed between *Aedes* and *Anopheles*. Collectively, these experiments demonstrate that the 3D-Feedy matches or exceeds the performance of the Hemotek system across two mosquito species, delivering higher fecundity in both cases while maintaining comparable egg viability.

## Discussion

In this study, we introduce and validate the 3D-Feedy membrane feeder, a low-cost, open-hardware device designed to improve the accessibility of mosquito blood feeding. Tested across two major disease vectors from different genera, *Anopheles gambiae* and *Aedes albopictus*, the device performed comparably to, or better than, the widely used Hemotek system. Feeding success was maintained at equivalent levels, and in both species, 3D-Feedy-fed females produced significantly more eggs without any reduction in egg viability.

Commercial systems such as Hemotek and Rutledge membrane blood feeders have become standard equipment for mosquito research laboratories. They are reliable and well validated, but their accessibility is constrained by several factors: they are expensive and tied to specialized suppliers, making them difficult to obtain or repair in many regions. Their reliance on tubing or cables connected to circulating water baths or bench-top power supplies fixes them in place, requiring cages to be carried to “feeding stations” rather than bringing feeders to the cages, whether located in different rooms, or incubators, or distant shelves. Capacity is another limitation: even the largest multi-unit Hemotek stations accommodate only six cages side by side. In large insectaries where dozens of cages may be fed daily, throughput can quickly become restricted, and expanding capacity demands additional “feeding stations”, an investment that consumes both money and valuable insectary space. These systems are also impractical for field settings.

Unsurprisingly, researchers have sought alternatives for decades, motivated by the need for tools that are affordable, user-friendly, easy to maintain, and ideally portable. Passive-heating designs, such as membranes coupled to pre-warmed vessels or metal blocks, are among the simplest and least expensive solutions (Tseng, 2003; Costa-da-Silva *et al*., 2013; Finlayson, Saingamsook and Somboon, 2015; Siria *et al*., 2018; Sri-in *et al*., 2020). Yet, because they lack active temperature regulation, they cool quickly and require repeated reheating, rendering them impractical for routine use. Low-cost electronics, 3D printing, and the DIYbio movement spurred a new generation of feeders. These designs reduce reliance on specialized glassware and demostrate that membrane feeders can be built with accessible components and basic skills. The SLAM feeder described by Dewsnup et al. is a notable example: an open-hardware device that demonstrated the feasibility of electrically heated, DIY feeders (Dewsnup *et al*., 2023).

Our 3D-Feedy builds on this foundation but incorporates several key refinements. It uses a digital thermostat with feedback control to maintain precise and stable temperatures, a machined aluminum block and large feeding plate to provide consistent heating across a large surface, and standardized USB-C power for flexibility, including operation from common powerbanks. Compared to Hemotek, the 3D-Feedy achieves equal or superior biological performance while avoiding the expense, supplier dependence, and scalability limits of commercial systems.

An important advantage of the 3D-Feedy is its repairability and simplicity. Inevitably, devices that see heavy use will develop faults, but because users assemble the feeder themselves, they gain the familiarity needed to diagnose and repair faults quickly. In contrast, commercial units can be costly to service, rely on proprietary parts, and often require or recommend specialist technicians. This self-sufficiency, combined with its low cost, makes the 3D-Feedy an accessible and sustainable option for laboratories of varying scale.

### Applications and scalability

The 3D-Feedy is ideally suited for contexts where cost, flexibility, and throughput are priorities. For smaller laboratories or new groups establishing colonies, it provides a reliable and inexpensive alternative to commercial systems. In teaching or community-engagement environments, the device’s simplicity makes it an effective tool for training in mosquito biology. In large insectaries, scalability is a major advantage: rather than relying on a single costly multi-unit station, researchers can build multiple 3D-Feedies at modest cost. Deploying multiple units allows parallel feeding across rooms, incubators, or experimental setups, greatly increasing throughput. Finally, its compatibility with portable USB-C power sources makes the 3D-Feedy practical in field sites and in facilities, where conventional feeders are impractical. Together, these features position 3D-Feedies as a robust, low-cost, and flexible option for mosquito feeding across a wide range of laboratory and field applications.

## Materials and Methods

### Design and construction of 3D-Feedies Parts list

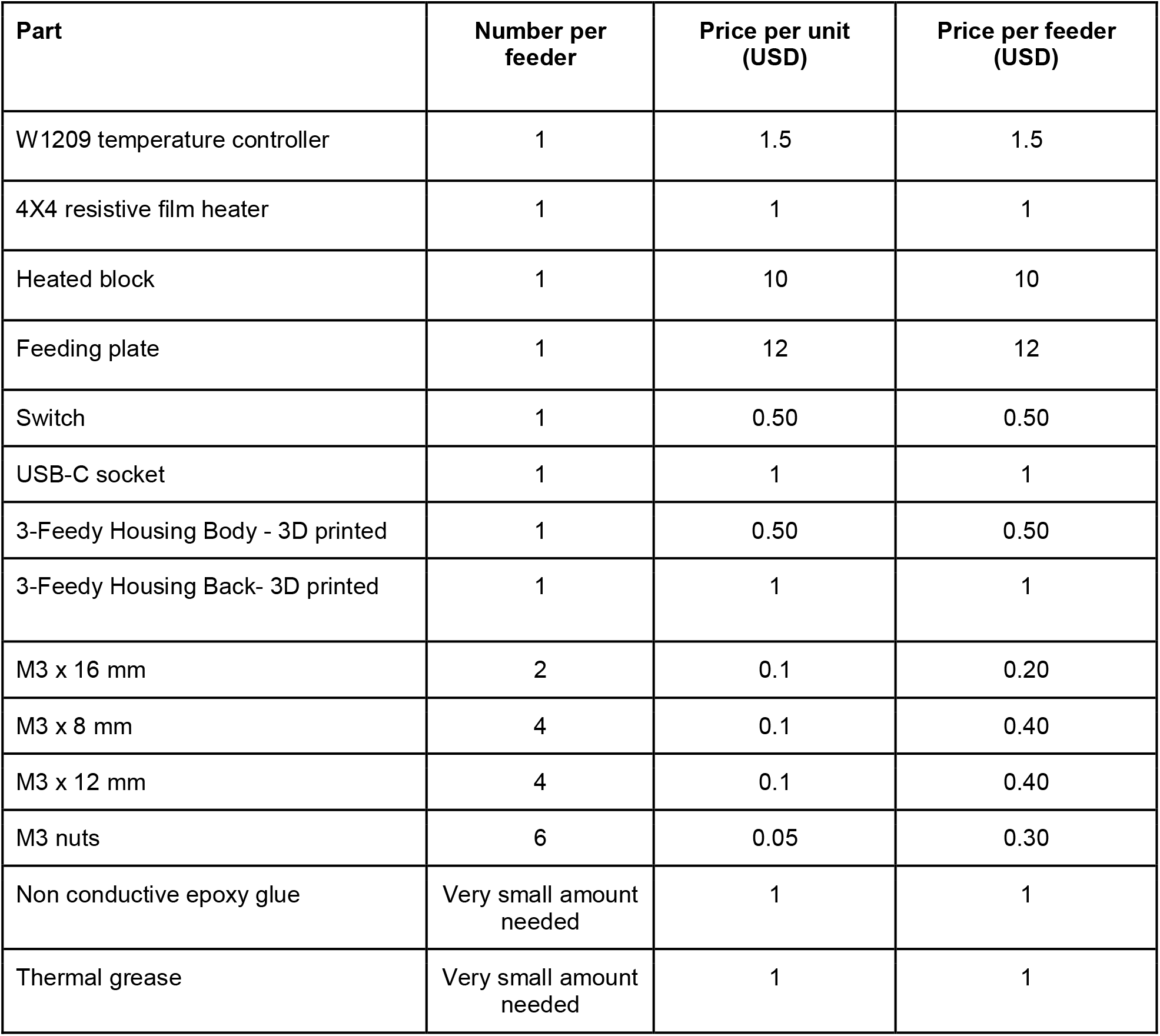

Parts list:
| Part | Number per feeder | Price per unit (USD) | Price per feeder (USD) |
| --- | --- | --- | --- |
| W1209 temperature controller | 1 | 1.5 | 1.5 |
| 4X4 resistive film heater | 1 | 1 | 1 |
| Heated block | 1 | 10 | 10 |
| Feeding plate | 1 | 12 | 12 |
| Switch | 1 | 0.50 | 0.50 |
| USB-C socket | 1 | 1 | 1 |
| 3-Feedy Housing Body - 3D printed | 1 | 0.50 | 0.50 |
| 3-Feedy Housing Back- 3D printed | 1 | 1 | 1 |
| M3 x 16 mm | 2 | 0.1 | 0.20 |
| M3 x 8 mm | 4 | 0.1 | 0.40 |
| M3 x 12 mm | 4 | 0.1 | 0.40 |
| M3 nuts | 6 | 0.05 | 0.30 |
| Non conductive epoxy glue | Very small amount needed | 1 | 1 |
| Thermal grease | Very small amount needed | 1 | 1 |

### Mosquito strains and rearing

*Anopheles gambiae* (G3 strain) and *Aedes albopictus* (FPA wild-type strain) mosquitoes were used in this study. Mosquitoes were maintained under standard insectary conditions (28 °C, 80% relative humidity, 12:12 h light:dark photoperiod) and reared according to established protocols.

### Mosquito blood feeding

Experimental females were three days post-emergence, mated, and unfed at the time of blood feeding. To minimize variation between groups, females for each replicate were derived from the same egg batch. Each treatment group consisted of three replicate cages containing 50 females per cage, giving a total of 12 cages (2 species × 2 feeder types × 3 replicates).

Blood used in all experiments originated from a single batch of thawed bovine blood, stored at – 20 °C. For *Aedes albopictus*, adenosine triphosphate (ATP) was added to a final concentration of 5 mg/mL to stimulate feeding, whereas no additive was used for *An. gambiae*. All feeders were fitted with Parafilm membranes of identical thickness and loaded with 2 mL of blood. Feeders were placed on top of mosquito cages and feeding was conducted in a 25 °C insectary working room rather than inside the environmental rearing chamber, to accommodate the Hemotek power supply. Each trial lasted 30 minutes. To measure feeding rates, females from each cage were aspirated, cold-anesthetized and the number of visibly blood-fed females was recorded for each cage.

### Mosquito egg production and laying

From each of the 12 cages, three groups of five engorged females were transferred to oviposition cages to quantify egg production (9 groups per treatment, 36 eggbowls in total). Egg bowls were introduced 48 h post-feeding and removed 24 h later to photograph the egg-containing filter paper, and eggs were counted using ClickMaster2000 (Eurac Biomedical Research, 2020). Mean egg output per female was calculated for each replicate.

### Larval Hatching

Eggs were incubated under species-specific protocols: For *An. gambiae*, eggs were allowed to mature in the eggbowl in standard insectary rearing water. For *Aedes albopictus*, hatching was stimulated using overnight incubated deoxidized hatching solution (Bellini *et al*., 2018), consisting of 0.25 mg nutrient broth (Merck-Sigma Aldrich, USA) and 0.05 mg baking yeast dissolved in 750 ml of distilled water, for two approximately two hours after which additional water was added to reach a total depth of 2 cm in the rearing container, followed by the addition of larval food.

### Statistical Analysis

Data were analyzed separately for *Ae. albopictus* and *An. gambiae*. Data normality was assessed using the Shapiro–Wilk test, and homogeneity of variances was examined with

Levene’s test. When both assumptions were satisfied, comparisons were made using two-sample *t*-tests; otherwise, non-parametric Wilcoxon rank-sum tests were applied. A two-way ANOVA was used to test for the effects of the feeding system, species, and their interaction on feeding rate and egg production. All tests were two-tailed with a significance threshold of _α_ = 0.05. Statistical analyses were performed in R version 4.4.1 (R Core Team, 2024) using the packages dplyr, car, and broom.

## Acknowledgements

We would like to thank Or Toren, Sigal Popovsky for their technical support and critical feedback of the manuscript. We are grateful to Nikolai Windbichler for insightful comments and for proposing the name 3D-Feedy.

## Author Contributions

ZG, AL and EO designed and built the 3D-Feedy. JD performed the initial testing and optimization of the feeders. GE performed all mosquito experiments and collected the data. ARL and PAP generated the figures. DSYZ and PAP analyzed the data and performed statistical analysis. EO and PAP secured funding. PAP wrote the manuscript with help from all authors. All authors read and approved the final manuscript.

## Ethics

All animals were handled in accordance and under the supervision of the ARO Institutional Animal Care and Use Committee approval number 2307-118-2-VOL-IL. All insect work was performed in HUJI facilities maintaining Arthropod Containment Level 2. This work received Institutional Approval and relevant authorizations from the Israel Ministry of Environmental Protection and Ministry of Agriculture (#31/2019).

## Supplementary Files 1–6

Design files for the 3D-Feedy housing for 3D-printing and aluminum plates for machining.

