## Supplementary figures and images for "3D-Feedy: An open and portable membrane feeder for mosquito research"

### Feeding_Plate_dimensions.png

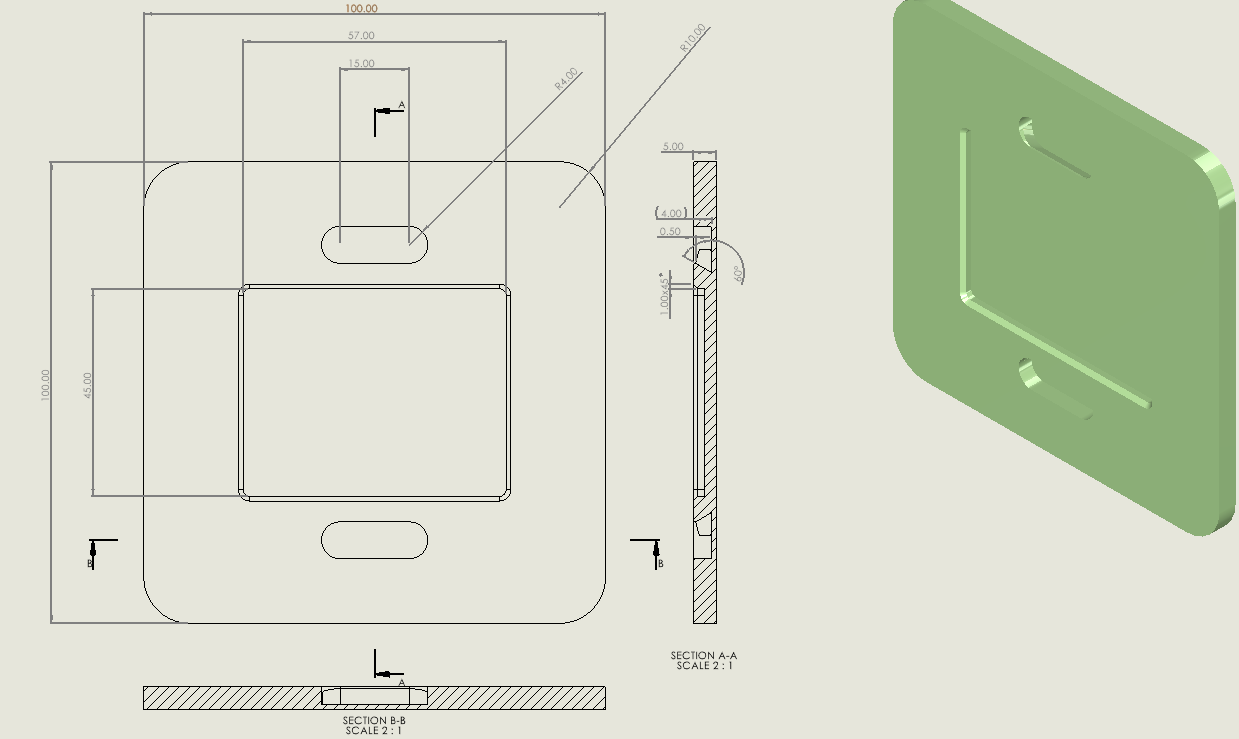

### Heated_Block_dimensions.png

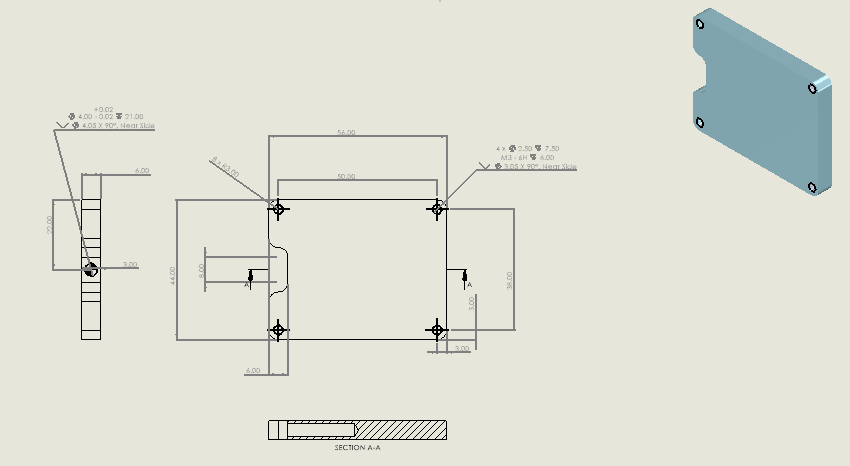
